# *mobileRNA*: a tool for efficient analysis of mobile RNA molecules in multiple genomes

**DOI:** 10.1101/2024.08.19.608270

**Authors:** Katie Jeynes-Cupper, Marco Catoni

## Abstract

In plants, mobile RNA molecules play a significant role in long distance signalling, with evidence of transport mechanisms and functional roles for both mobile messenger RNA (mRNA) and small RNA (sRNA) molecules. The movement of RNAs between distant tissues can be revealed in plant grafting experiments involving different genotypes (hetero-grafting) associated to genomic analysis, using the natural polymorphisms between the genotypes to discriminate between RNAs produced in the sampled tissue from those which have travelled from distant organs. However, the hight level of noise in the analyses of RNA sequencing datasets involving two different genotypes, and the lack of a standardised pipeline for the analysis of grafted plants, make the identification of natural mobile RNA molecules a challenge. Here, we introduce a pipeline integrated into an R package, *mobileRNA*, which performs simultaneous alignment of RNA sequencing samples on a merged reference genome. Using simulated datasets, we show that *mobileRNA* can identify putative mobile RNAs with unprecedented efficiency in absence of post-processing steps.

## 1. Introduction

Plant grafting is a natural process consisting in joining two plants into one, which continue to grow in a chimeric condition. In horticulture, artificial grafting has been historically employed to introduce or improve desirable agronomic traits in plants species of economical relevance but has also emerged as a successful research tool in scientific discovery (Jeynes-Cupper and Catoni 2023). In this practise, the plant which contributes to the shoot is defined as the scion and the part that contributes to the lower stem and roots is referred to as the rootstock. Due to the systemic connection established in successful grafting between the scion and the rootstock, plant grafting has been used to study the roles of mobile proteins, RNAs and hormones in the plant signalling pathway. In recent years, researchers have used grafting to demonstrate RNA transport between the shoots and roots of specific molecules, and their implications to genetic and epigenetic regulation (Molnar, Melnyk et al. 2010, Mahajan, Bhogale et al. 2012, Notaguchi, Higashiyama et al. 2015, Lewsey, Hardcastle et al. 2016, Li, Wang et al. 2021, Rubio, Stammitti et al. 2022).

To systematically study the mobile RNA populations within a plant model, high throughput genomic analysis with Next-Generation Sequencing (NGS) technologies can be used to provide quantitative insights and genome-wide variations between conditions (Deshpande, Chhugani et al. 2023). For the investigation of mobile RNA, RNA sequencing (generally known as RNAseq or mRNAseq), is the method of choice to quantify gene expression, while the sequencing of non-coding small RNAs (sRNAseq) with regulatory roles can be used to quantify their accumulation in plant tissues (Li 2018). These approaches are based on the extraction of RNA from a sample, the conversion to complementary DNA (cDNA) and the generation of libraries to be processed into NGS. If a reference genome assembly is available for the studied organism, the resulting sequenced reads are then aligned to it, and the RNA expression levels can be analysed (Deshpande, Chhugani et al. 2023). Alignment tools have been optimised to balance speed and accuracy, and the most advanced examples consider the presence of errors occurring during sequencing and reverse transcription, as well as the presence of potential structural variants, like single nucleotide polymorphisms (SNPs), which all generate variation between the sequenced reads and the reference genome (Kim 2019). These approaches take as assumption that all or most sequenced nucleic acid are assembled in the reference genome and determines the best possible location to place a sequenced read in the given genome, in accordance with the alignment quality thresholds defined by the chosen parameters. While this is optimal for a standard RNAseq experiment where a single genotype is considered, the alignment output become less predictable in instances in which biological samples contain genetic sequences from multiple organisms and where both are investigated, as in the case of heterografted plant combinations.

In past analysis, when samples obtained from heterografted plants has been aligned to a reference, researchers have utilised different approaches to use genetic polymorphisms to discriminate the origin of the RNA molecules (Notaguchi, Higashiyama et al. 2015, Li, Wang et al. 2021, Rubio, Stammitti et al. 2022). These pipelines frequently use independent single-reference mapping approaches and recover un-mapped reads (Notaguchi, Higashiyama et al. 2015, Xia, Zheng et al. 2018, Wang, Wang et al. 2020, Li, Wang et al. 2021, Rubio, Stammitti et al. 2022), or apply downstream variant call analysis and additional statistic to discriminate noise from genuine polymorphisms (Tomkins, Hoerbst et al. 2022). However, in both cases mis-aligned reads can originate by incomplete information in the mapping steps, which is always performed on a single reference.

Here, we introduce an alternative approach to standardise the analysis of long-distance RNA movement using plant heterografting. We implemented a simultaneous alignment of RNAseq datasets to two (or more) independent reference genomes, so that all possible origins of the RNA molecules are provided in a single alignment step.

Using simulated plant grafting experiments obtained from sequenced datasets, we demonstrate that our pipeline can map reads to their genome of origin accurately with a strong reduction of data noise, although the similarity of the two genomes used as references plays the main role in discriminating of the placed reads across the two genomes. To allow reproducibility and easy usage of our pipeline to general biologists, we implemented our approach into a R package for RNAseq and sRNAseq analysis, which we named *mobileRNA*, available in the Bioconductor repository (Jeynes-Cupper and Catoni 2023).

## 2. Methods & Materials

### Description of *mobileRNA* pipeline

We assembled an analysis pipeline to process and quantify the expression of graft-mobile RNAs for sRNAseq or RNAseq datasets with single or multiple genomes. When two or more genomes are used, these are merged to perform a single alignment step, rather than multiple mapping on single references as previously done.

The workflow is applied when samples are mapped on the merged references, and it aims to locate the expression of RNAs from more than one genome. For example, graft-mobile RNAs traveling from the rootstock-to-scion in an inter-specific heterograft. The input dataset of the pipeline consist in the cleaned sequencing reads from sRNA sequencing or mRNA sequencing in FASTQ format, from control and treatment samples (if available), along with the single or multiple genome assemblies (FASTA), together with the corresponding genome annotations files (GFF), if available (Supplementary Figure 1 for further detail). The full procedure is available in the Bioconductor/R package *mobileRNA* (available at https://bioconductor.org/packages/mobileRNA).

### Merging of reference genomes and annotations, alignment and raw count

To discriminate the genomic origin of sequencing reads in multiple genomes during the aligning step, *mobileRNA* generated a new merged reference genome (in FASTA format) to be utilises in the alignment of reads. A unique prefix is added to each pseudomolecule (chromosome/contig/scaffold) in the genomes to maintain them distinguishable in the downstream analysis. Similarly, a single annotation is generated by merging the original annotation files of the single genomes. The genome references and annotations were collated from public repositories and described in Supplementary Table 1.

If mRNA reads are used as input, the alignment step is performed with HISAT (default settings) (Kim, Langmead et al. 2015), keeping uniquely mapped reads, followed by raw count performed for each gene with HTSeq (Anders, Pyl et al. 2015) (Figure 1). If sRNA reads are used as input, they are aligned to the merged reference file using Bowtie (Langmead, Trapnell et al. 2009), keeping uniquely mapped reads, and ShortStack (Axtell 2013) is used to undertake sRNA cluster analysis. For the analysis of sRNAs, ShortStack is first run once to generate a sRNA cluster annotation for each sample. Then, each sample annotation is merged into a single file in R using the GenomicRange package (Lawrence, Huber et al. 2013), and ShortStack is run a second time to quantify the coverage of each cluster.

**Figure 1.**
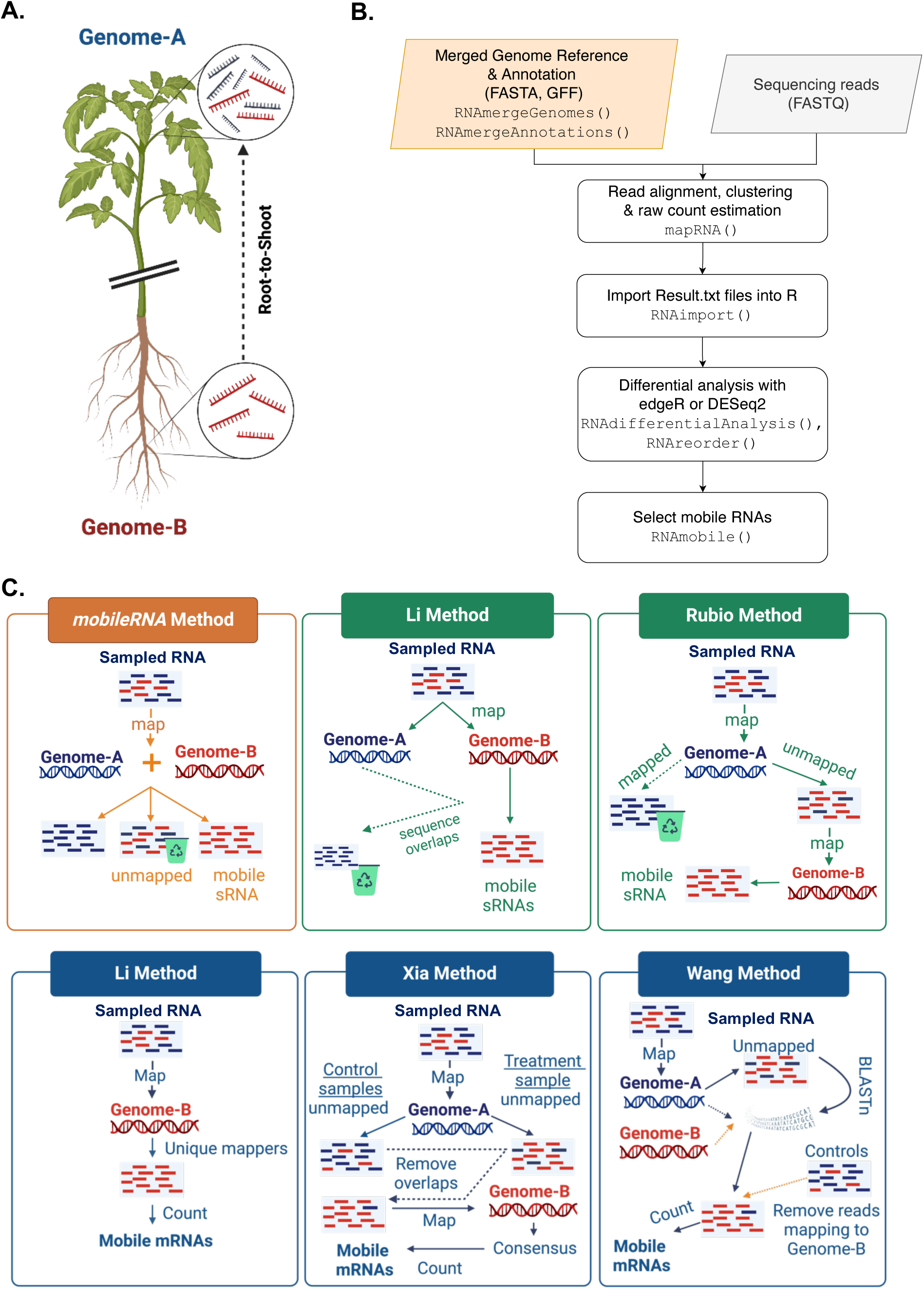
*mobileNA* workflow and implemented bioinformatic pipelines. A) Diagram of a grafting experiment involving mobile RNA molecules (movement assumed in the root-to-shoot direction). The sampled material (assumed as scion) will include molecules produced by the genome A (in blue) as well as mobile molecules that have travelled from the Genome B (in red). B) The simplified *mobileRNA* workflow implemented in R (see Supplementary Figure 1 for a more detailed representation). The genome references for the scion and rootstock (Genome Reference A and B) are merged into a single genome reference where sequences (e.g. chromosomes) from each genome remain distinguishable. The sequenced reads, either from sRNAseq or mRNAseq, are aligned to the merged reference genome, followed by clustering (only for sRNAseq) and raw quantification. The results are saved to the users desired directory, and then imported into R for downstream analysis, consisting in the selection of the putative graft-mobile RNA candidates. A full description of additional resources and the full source code of *mobileRNA* pipeline is available on Bioconductor repository (available at https://bioconductor.org/packages/mobileRNA). C) Schematic diagrams of the mobileRNA pipeline and the re-implemented pipelines utilised for benchmarking, including Li and Rubio methods (Li, Wang et al. 2021, Rubio, Stammitti et al. 2022) (for sRNA analysis), and the Li, Xia and Wang methods (for mRNA analysis) (Xia, Zheng et al. 2018, Wang, Wang et al. 2020, Li, Wang et al. 2021). Blue and red lines represent reads from the Genome A (blue) and Genome B (red), respectively. See methods section for more detail related to each approach.

### Genome mappability and index calculation

The genome mappability estimates the degree of repetitiveness of a genome (Derrien et al. 2012), but can also be used to evaluate how unique or repetitive a DNA regions is in a given genome (Catoni et al. 2017). Mappability is expressed, at every genome location, with a score from 0 to 1, where 1 is represents a unique region and 0 represents a hypothetical infinitely repeated region. Here, we used GenMap (1.3.0) (Pockrandt, Alzamel et al. 2020) to compute the mappability at single nucleotide resolution (-K 30) for each of the genome used, taken alone or in combination in pairs (merged genomes). The mean mappability value across each exonic region was determined using BEDtools (bedtools map -c 4 -o mean) (Quinlan and Hall 2010). For each gene, a single mean was obtained by averaging the value of each exon weighted on they nucleotide length. These averaged values have been used to estimate gene repetitiveness in single or merged genomes. To compare the mappability of the different merged genome combination, we defined a *genome mappability index* as the proportion of exonic regions that is above a mappability threshold of 0.5.

### Quantitative analysis and differential expression

Raw counts of either genes (mRNA analysis) or sRNA clusters are imported into R and organised into a data frame object. Quantitative data, including Fragments Per Kilobase of transcript per Million (FPKM) for genes, or Reads Per Million (RPM) in each sRNA cluster, are automatically calculated. If the experimental design includes control and treatment conditions (e.g. self-grafted control plants and hetero-grafted plants), differential expression analysis can be performed with either *edgeR* (Robinson, McCarthy, & Smyth, 2010) or *DESeq2* (Love, Huber, & Anders, 2014) R packages, which are both integrated into the *mobileRNA* pipeline. For either statistical test, the *P*-values are adjusted for multiple testing using Benjamini and Hochberg’s method (Benjamini Y, 1995) to control the false discovery rate (FDR).

### Dicer call

When sRNAseq samples are analysed, for each sRNA cluster identified ShortStack attempts to infer the putative Dicer enzyme originating the sRNAs (dicercall). This can differ among samples and can have biological significance (Shahid and Axtell 2014). *mobileRNA* uses this information to predict a “consensus” dicercall class for each cluster. This is done by comparing the dicercall for all clusters in each sample or specific samples, and then selects for the most abundant call. In cases of a tied values, the dicercall is set unclassified (but a valid dicercall value is prioritised over an unclassified result in a tie between the two).

### Mobile RNA analysis

To detect putative mobile RNAs, *mobileRNA* uses the raw count of features (either for mRNA or sRNAs) obtained from the alignment performed using the merged genomes. In this way, the mapping tool algorithm discriminate for each read the best placement on the two genomes accordingly to the alignment parameters set by the user. The gene annotated features (for RNAseq) or the sRNA clusters aligned to the distant genome reference (Genome B) containing reads are assumed to represent graft-mobile RNAs. Other filtering steps can be applied, as following:

1) If control self-grafted samples are available (grafting experiment using only Genome A genotype), these are used to select the graft-mobile RNAs as these molecules should exhibit an absence of read counts in control samples (assuming that their alignment on Genome B is due to incorrect mapping).
2) If biological replicates are available, it can be applied a threshold that considers the number of hetero-grafted replicates which contained reads.
3) Candidates can be further filtered by a minimal value of coverage observed at Genome B reference, however, this is not used as default as graft-mobile RNAs candidates might exhibit relatively low read counts compared to RNA population associated to the genotype of the sample of origin.

### Output results

Upon completing the RNA analysis, *mobileRNA* generates a data frame that serves as a structured repository of information with rows representing the sRNA clusters or mRNA transcripts. Additional information provided includes, for the sRNA analysis: the dicercall for each replicate, consensus dicercall, the number of replicates contributing to the consensus dicercall, count mean, log fold change (compared to control condition, if available), p-value, adjusted p-value, RNA sequence, consensus RNA sequence and RPM. For the mRNA transcripts, the final output includes the genomic location, pseudomolecule, gene name, the number of replicates contributing to a read count, count mean, log fold change (compared to control condition, if available), p-value, adjusted p-value, and FPKM. In the core RNA analysis output, the above information is provided for all the molecules analysed in the reference (either genes or sRNA clusters). If the mobile RNA analysis is performed, only the filtered graft-mobile RNAs are provided in the output

### Visualising data & additional features for assisting functional analysis

We also implemented in *mobileRNA* package additional exploratory data analysis for quality control purposes which uses multidimensional scaling to illustrate the relationships between samples. This includes principal component analysis (PCA), which is performed using the R package *DESeq2* (Love, Huber, & Anders, 2014), an analysis of the distribution of the sRNA dicercall classes within a sample or the consensus across the experimental design. In addition, the RNA expression can be visualised with a heatmap using hierarchical clustering of normalized RPM/FPKM values transformed logarithmically, commonly referred to as “rlog” values. Similarly, users can plot the distribution of genomic features overlapping sRNA clusters with the assistance of genome annotation files (GFF). A full description of additional resources and the full source code of *mobileRNA* pipeline is available on Bioconductor repository (available at https://bioconductor.org/packages/mobileRNA).

### Re-implementation of known pipelines for graft-mobile RNA identification

For benchmarking purpose, we re-implemented previous methodologies utilised in published research papers aiming to identify the mobilisation of mRNAs or sRNAs in plant inter-specific heterografts. In a basic comparison, these methods commonly aligned reads to each genome separately, followed by post-alignment screening of genetic variants (Figure 1C).

For the analysis of graft-mobile sRNAs, we re-implement two previous pipelines. For simplicity, each method has been named after the first author’s name; known as Li method (Li, Wang et al. 2021) and Rubio method (Rubio, Stammitti et al. 2022). The Li method was taken from a study which grafted common bean (*Phaseolus vulgaris*) and soybean (*Glycine max*) but was limited to one replicate per condition (Li, Wang et al. 2021). The Li method aligns a sample to each genome (sampled genome and distant genome) separately and then selects reads which were aligned to the distant genome but were unmapped to the sampled genome. The Rubio method was taken from a study utilising grapevine species; Cabernet Sauvignon grapevine (*Vitis vinifera*) and Gloire de Montpellier grapevine (*Vitis riparia*) (Rubio, Stammitti et al. 2022). Sample reads are first mapped to the tissue sampled genome (Genome A), then all unmapped reads are retrieved and re-mapped to the distant (Genome B) (Figure 1C).

For the analysis of graft-mobile mRNAs, the *mobileRNA* approach was compared to three previous pipelines. These methods were named Li-mRNA method (Li, Wang et al. 2021), Xia method (Xia, Zheng et al. 2018), and Wang method (Wang, Wang et al. 2020) (Figure 1C).

The Li-mRNA method (Li, Wang et al. 2021) aligns cleaned reads to the distant genome using STAR (Dobin, Davis et al. 2013) and only mRNA features with uniquely mapped reads were extracted and their normalised abundance estimated with CPM (counts per million) using BEDtools (Quinlan and Hall 2010) (Figure 1C).

The Xia method (Xia, Zheng et al. 2018) aligns cleaned reads to the sampled genome origin using HISAT (Kim, Langmead et al. 2015) with default settings. The unmapped reads are retrieved using SAMtools (view -f 4) from both heterografts samples and self-grafted control (Danecek, Bonfield et al. 2021). Then reads from the hetrografted samples which perfect matches with the self-grafted control are discarded using BEDtools (intersect -wa -wb) (Quinlan and Hall 2010). The remaining unmapped reads are then mapped to the distant genome using HISAT (Kim, Langmead et al. 2015). Then only aligned reads that match between at least two of three samples were considered associated to mobile mRNAs. These reads are then re-mapped to the distant genome and raw counts of associated transcript is calculated using HTSeq (-- order=pos -a=0 --stranded=no --mode=union --nonunique=all --type=mRNA) (Anders, Pyl et al. 2015) (Figure 1C).

Finally, the Wang method (Wang, Wang et al. 2020) aligns clean reads to the sampled genome using Bowtie2 (sensitive-local mode: -D 15 -R 2 -N 0 -L 20 -i S, 1, 0.75) (Langmead and Salzberg 2012), and extracts the unmapped reads using SAMtools (view -f 4) (Danecek, Bonfield et al. 2021). For each sample, the unmapped reads were blasted against both the sampled and distant genomes, independently, using BLASTn (Altschul, Gish et al. 1990) (-max_target_seqs 3 -e value 1e-5 -outfmt 6 -num_threads 8). Reads from self-graft controls which aligned to the foreign genome were considered as false positives and were subtracted from the pool of potential mobile mRNAs. The remaining reads were used to obtained raw count of associated transcripts with HTSeq (Anders, Pyl et al. 2015), and considered potentially mobile (Figure 1C).

### Generating simulated datasets

To generate the sRNA blended data, we retrieved real RNAseq or sRNAseq datasets obtained from tomato, eggplant and Arabidopsis (Col-0, Ler-0 and C24 accessions) samples growing in normal conditions, downloaded from the public Sequence Read Database (SRA) repository. We first paired samples to simulate hetero-grafting combinations of eggplant and tomato, or among different *Arabidopsis thaliana* accessions (Col-0, C24 and Ler-0), as detailed in Supplementary Table 2.

Then, to generate blended sRNA datasets, we first perform classic analyses (mapping on a single reference) of the real sequenced samples chosen to represent the genome (Supplementary Table 1 & Supplementary Table 2). The raw reads were trimmed using trimmomatic (Bolger, Lohse et al. 2014) and then uniquely aligned to the reference of the distant genome using Bowtie (bowtie -p 6 -v 0 -a -m 1) (Langmead, Trapnell et al. 2009). Depending by the pair combination, reads were aligned to the *Solanum lycopersicon* (Tomato) Version 4 assembly (Hosmani, Flores-Gonzalez et al. 2019) taken from the Sol Genomics Network (https://solgenomics.net/), the *A. thaliana* Landsberg (Ler-0) de novo assembly version 2 (Jiao and Schneeberger 2019) taken from “The Arabidopsis Information Resource” (https://www.arabidopsis.org/), or *the A. thaliana* C24 de novo assembly version 2 (Jiao and Schneeberger 2019) taken from “The Arabidopsis Information Resource” (https://www.arabidopsis.org/). Then, for each individual sample the alignment file was passed to ShortStack (V4.0.2) (Axtell 2013) which undertook sRNA cluster analysis (--threads 6 --pad 200 --mincov 0.5 --nohp). We filter all detected cluster to have a single predicted dicercall, a non-zero raw count and the RNA sequence to not align falsely to the reference chosen as sampled genome. We then selected randomly 150 cluster loci and we retrieved all reads associated to them from the original alignment file using SAMtools (view -b -L) (Danecek, Bonfield et al. 2021), and converted from BAM to FASTQ format using BEDtools (bamtofastq) (Quinlan and Hall 2010). Then, we used the obtained reads to spike real sRNA datasets obtained from samples of the species used to simulate the sampled genome (Supplementary Table 5). The sets of sRNAseq blended datasets generated with this method was named as “Random-Format”. To have more control on the expression level of the simulated mobile sRNAs, we also generated blended datasets selecting 150 cluster loci in three groups of 50 loci each associated to respectively a low (< 5x mean quartile), average (mean) or high (> ∼5x mean quartile) read coverage in the original alignment. This second approach was named “Tier-format” (Supplementary Table 3, Supplementary Table 5).

To generate simulated mRNA data, we used the R package *polyester* (Frazee, Jaffe et al. 2024) to produce sets of synthetic raw RNAseq reads from 150 genes selected randomly from the annotation of the reference used as the distant genome (genome B) (Supplementary Table 2). The baseline mean number of reads per transcript was calculated based on transcript length, a coverage of 20 and 100 bp read length. The simulated expression for each of the 150 transcripts was set at three different values (1, 5, or 10) distributed equally in three groups of transcripts (composed by 50 transcript each). To generate blended datasets which simulate heterografting, we spiked 1000, 5000, 10,000 or 50,000 of these synthetic reads into real RNAseq datasets sequenced from biological samples of the species chosen as the sampled genome (Supplementary Table 4). Therefore, for each heterografted combination tested, we generated blended datasets with different amounts of reads from putative mobile RNAs, spanning from approximately 0.006% (for spiking with 1000 reads) to 0.293% (for spiking with 50,000 reads) of the total sequenced pair reads (in average between 3-13 million). The same real datasets used for generating the blended data, without any reads spiking, where used to simulate self-grafted control samples.

Within each blended mRNA and sRNA dataset, the distribution of reads in each library were comparable (Supplementary Figure 2, Supplementary Table 4). In both instances, the Col-0/Ler-0 and Col-0/C24 simulated datasets were more similar in read distribution in comparison to the Eggplant/Tomato simulated dataset where they demonstrated mRNA population with a greater mean in FPKM (Supplementary Figure 2, Supplementary Table 4). The mappability of the spiked simulated graft-mobile mRNAs were assessed in the genome of their origin and within the merged genome. The merged genome contains the references for the genome of origin and the genome representing the genotype of the destination tissue. It was expected that within the merged genome, the mRNA transcripts would have a reduced mappability (i.e. less unique), and especially in the Col-0/Ler-0 simulated dataset where the genomes are highly similar and separated by several SNPs within the coding regions. In general, this was demonstrated in the data where over 92% of mRNA transcripts showed a reduced or unchanged mappability.

## 3. Results

### Implementing a merged genome approach for efficient graft-mobile RNA identification

We assembled a complete pipeline for the efficient detection of mobile mRNA and sRNA molecules, starting from RNAseq and sRNAseq datasets obtained in plant grafting experiments involving two different genotypes (heterografts). Our approach assumes that polymorphisms (SNPs and structural variants) between the two genomes will allow the discrimination of RNA molecules produced in the distant genotype (e.g. rootstock) which travelled in the sampled genotype (e.g. scion) (Figure 1A and 1B). In previous studies with a similar design, the read aligning step was performed to only one genome (or to each genome separately) and it was followed by post-alignment screening of potential mobile molecules, requiring in some cases additional statistic to discriminate real polymorphisms from noise (Figure 1C) (Notaguchi, Higashiyama et al. 2015, Xia, Zheng et al. 2018, Wang, Wang et al. 2020, Li, Wang et al. 2021, Rubio, Stammitti et al. 2022). To avoid such post-alignment analysis and to optimise the identification of reads potentially originated from mobile RNAs, we implemented an alignment step performed with both involved genomes at the same time. This is obtained by merging the two reference genomes of the genotypes used in a grafting experiment into a single file, which is then taken as main reference for mapping (see methods) (Figure 1B). Then, uniquely mapped reads sequenced in the sampled genotype which have found aligned to the reference of the distant genotype are recovered and used to determine which annotated features (transcripts or sRNA clusters) have potentially moved into the sampled genotype (Figure 1B).

To test our procedure, we utilised combinations of genomes known to be graft compatible, merging the eggplant (*Solanum melongena*) and tomato (*Solanum lycopersicum*) references to simulate an interspecific grafting combination, and two combinations of *Arabidopsis thaliana* accessions for intraspecific grafts (Supplementary Table 1). For the latter, we paired the Col-0 reference with either C24 or Ler-0 accessions, for which *de novo* genome assemblies are available (Schneeberger, Ossowski et al. 2011, Jiao and Schneeberger 2020).

The merging of two genomes into a single reference will reduce the mapping efficiency of genomic sequencing reads, due to the presence of homology between the two genotypes involved. To estimate this effect, we calculated the mappability score for coding regions annotated in each merged genome in comparison to the two individual references involved (Figure 2).

**Figure 2.**
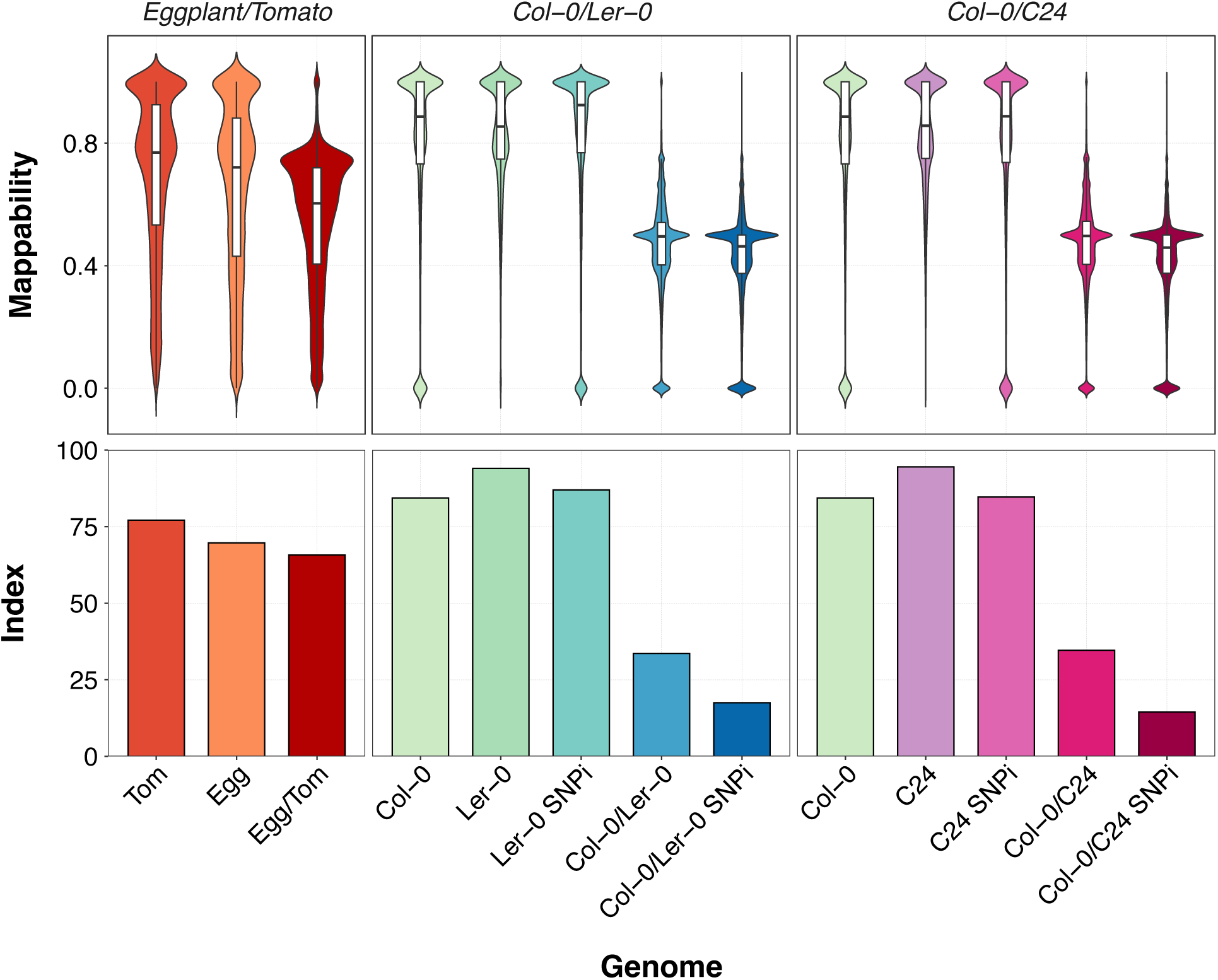
The genome mappability for individual and merged genome reference. The genomes and their origin are outlined in Supplementary Table 1. Some genome names have been abbreviated, including Tomato referred to as “Tom”, Eggplant referred to as “Egg”, and “SNPi” refers to a single nucleotide polymorphism (SNP) injected genome reference. The genome mappability value is represented from 0-1 where 0 represents a highly repetitive region and 1 signifies a highly unique region. The genome mappability is calculated across the genome, and the plot illustrated the distribution of mappability across coding regions of the genome. For each analysis, the mappability for the individual and merged genomes are illustrated as violin plots. The Index (%) is a measure of the proportion of the genome which is above a mappability score of 0.5 and is shown as a bar plot for each genome reference.

For the Ler-0 and the C24 Arabidopsis accessions, we calculated a mappability score using either the *de novo* assembled references, or the references obtained by injecting SNPs into the Col-0 reference. As expected, we observed a reduction in mappability in all combinations of merged genomes, compared to the corresponding individual references, with a drop of 15% in the average mappability for the Eggplant/Tomato combination and about 40% for all Arabidopsis combinations (Col-0/C24 and Col-0/Ler-0, independently using the SNP-injected or *de novo* assembled references). In the Eggplant/Tomato merged genome the proportion of coding regions with a mappability above 0.5 was 66% (compared to 70% and 77% in eggplant and tomato reference, respectively). In the Arabidopsis combinations built using the *de novo* assembled references, the calculated average mappability was 34% and 35%, respectively for the Col-0/Ler-0 and Col0/C24 merged references, values which dropped respectively to 18% and 14% in the combinations including SNP-injected references (Supplementary Table 1 and Figure 2). These results confirm that interspecific grafting provides better potential discrimination of RNA molecules compared to interspecific grafting, and indicate that, for Arabidopsis, *de novo* assembled references allowed to potentially map (mappability > 0.5) almost twice the number of coding regions compared to references obtained with SNP injection. Therefore, only *de novo* assembled references were used in following analyses (Figure 2).

### Benchmarking *mobileRNA* with blended RNA datasets

Simulated synthetic datasets are often used for benchmarking genomics pipelines, because they give control on variables which are normally unknown in biological data. Nonetheless, synthetic data are artificially constructed and often are not reflecting real biological contexts. Therefore, to benchmark *mobileRNA* performance we generated blended RNAseq datasets starting from real experimental data, obtained by spiking RNA reads into datasets from different genotypes to simulate the potential systemic movement of molecules in each tested heterografted plant combination (Lähnemann, Köster et al. 2020). This approach provides complete control of real positives and allows for benchmarking both false negative and false positive rates across different systems used to retrieve mobile RNA molecules. To generated datasets compatible with the real scenarios, we used data samples from published RNAseq and sRNAseq experiments performed with graft-compatible genotypes, including: i) Eggplant (*Solanum melongena*) and Tomato *(Solanum lycopersicum*), ii) *Arabidopsis thaliana* Columbia (Col-0) and Landsberg erecta (Ler-0) accessions and, iii) *A. thaliana* Col-0 and C24 accessions; always considering the first genotype as the tissue sampled genome (assumed as scion) and the second genotype as the distant genotype (assumed as rootstock). For these combinations, we built blended data for both sRNA and mRNA datasets by spiking reads from the sampled of the distant genome into the dataset of the sampled genome, simulating the movement of 150 sRNA (Supplementary Table 5) or mRNA (Supplementary Table 4) molecules at different level of expression and relative amount in the pool of sequenced reads (see method).

### Analysis of sRNA datasets

We used our sRNA blended datasets to benchmark the performance of *mobileRNA* in the analysis of mobile sRNA molecules, compared with two previously established approaches based on alignment steps performed on single genomes, namely the Li (sRNA) and the Rubio methods (Figure 1C).

To determine the effectiveness of each analytical method, the number of correctly identified mobile sRNAs clusters (true positives) were recorded along with the number of incorrectly identified mobile sRNAs (false positives) and, of the spiked sRNA cluster, the number of those which were not identified (false negatives) (Figure 3A). Of 150 simulated mobile sRNA clusters, *mobileRNA* recovered ∼97%, 60% and 24% respectively in the Eggplant/Tomato, Col-0/C24 and Col-0/Ler-0 combinations (Supplementary Table 7). The performance inversely correlates with the genetic similarity between the two genotypes used in the simulated experiment (Schneeberger, Ossowski et al. 2011, Jiao and Schneeberger 2020).

**Figure 3.**
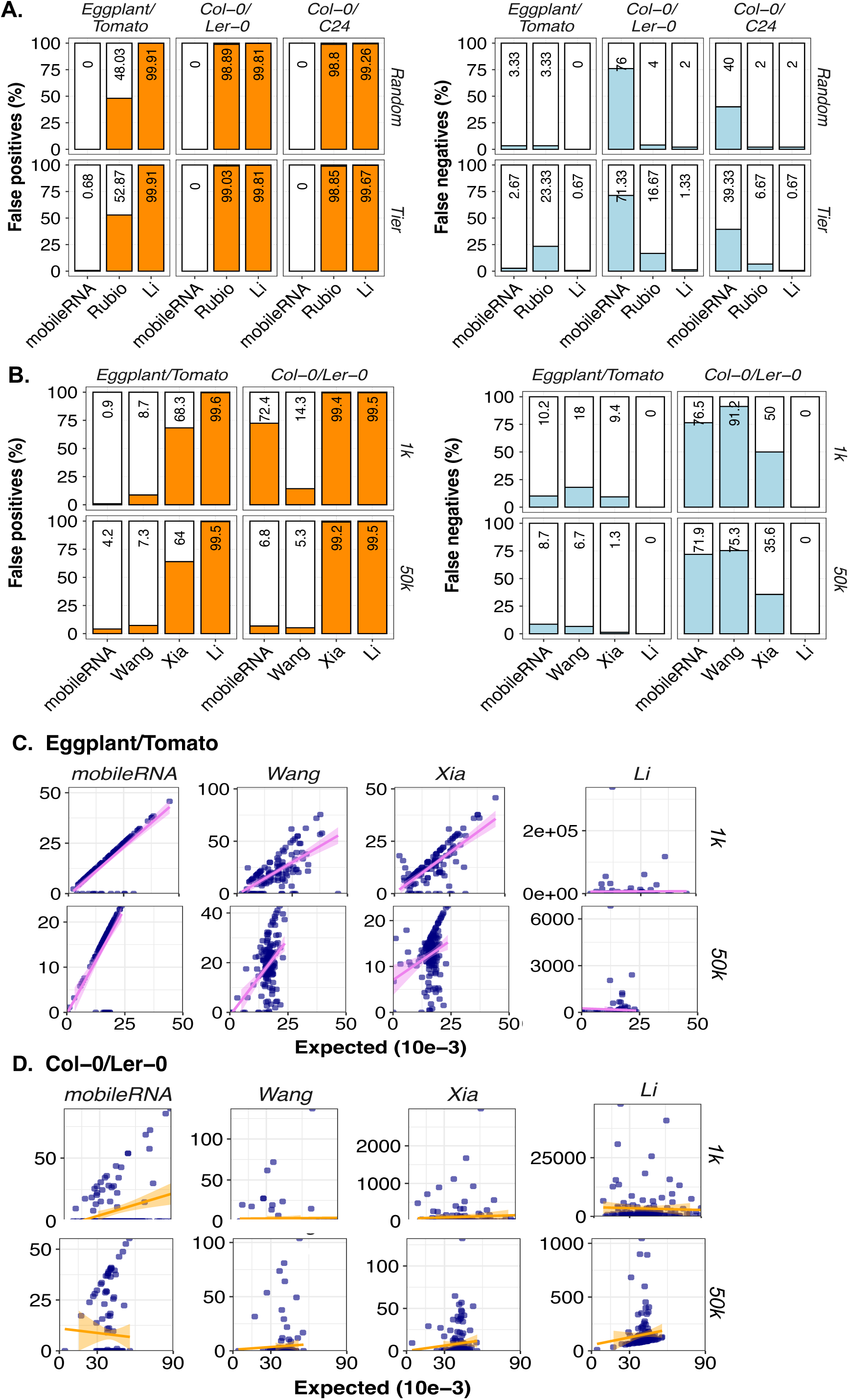
Comparative analysis of mobile RNAs identified by the analytical methods in each simulated dataset. A) For each sRNA simulated experiment, proportion of false positives on the total number of clusters identified as mobile, and proportion of false negative (expressed as percentage of the total spiked clusters) for each methodology tested. B) For each mRNA simulated experiment, proportion of false positives on the total number of transcripts identified as mobile, and proportion of false negative (expressed as percentage of the total spiked transcripts) for each methodology tested. Note that only spiking with 1K and 50K are displayed (5K and 10K are available as Supplementary Figure 5). C). Scatter plots displaying, for each mRNA methodology applied, the correlation of the expression measured (as FPKM) compared to the expected value (obtained by mapping on a single reference the samples from where the spiked reads where obtained). Data are show for both the Eggplant/Tomato dataset and the Col-0/Ler-0 dataset (data from 5K and 10K are available as Supplementary Figure 5.

Moreover, when we used simulated mobile sRNA clusters at a specific expression level (Tier-Format, see method), we observed that the performance of the *mobileRNA* method is mostly not affected by the level of expression (Supplementary Figure 4). The lower performances of the Li and Rubio methods in term of false negative rates appear mostly toward the lowly expressed sRNA clusters (Supplementary Figure 4) but, contrary to *mobileRNA*, these methods are not significantly affected by the degree of similarity of the two genotypes used in the simulation. These results, taken together, indicate that the recovery rate of mobile sRNA is affected by their abundance, irrespective by the method used. However, if a merged genome mapping approach is used, the genetic similarity of the genotype involved play also an important role.

Then the false positive rate was estimated, *mobileRNA* displayed remarkably outstanding performance in all simulations, recovering less than 1% of false positives independently by the genotype combination or the level of sRNA expression. By contrast, Li and Rubio methods retrieves respectively 99% and ∼48% of false positives in the Eggplant/Tomato combination, independently by the level of sRNA expression, while both method retrieves over 98% of false positives in the Arabidopsis simulations. This result indicates that the alignment to merged genomes can strongly improve the sensitivity of mobile sRNA detection reducing the background noise (Figure 3C, Supplementary Table 7).

### Analysis of mRNA datasets

Compared to sRNAs, mobile mRNA are longer molecules that must be transcribed from genes in the nucleus and must be actively transported across cells compartment. For this reason, their accumulation in plant organs distant from their tissue of origin could be in many cases lower compared to the abundance of averaged expressed genes. To account for potential low proportion of mobile mRNAs, we generated blended datasets with 1k, 5k, 10k, and 50k reads, corresponding to 0.002% to 0.451% of a full library of RNAseq, using the un-blended sample as self-grafted control (Supplementary Table 4). Then, for each simulated data set, the total number of simulated mobile mRNAs were determined with *mobileRNA* (see methods) and compared with the re-implementation of three previous systems based on single genome mapping, including the Wang, Xia and Li methods (Figure 1C). For each pipeline, the number of identified mobile mRNAs transcripts (derived from the reads mapped uniquely on the distant genotype) were recorded along with false positive and false negative (Figure 3B-C, Supplementary Figure 5, Supplementary Table 6).

The proportion of retrieved false positive decreases fractionally as the spiked reads used increased from 1k to 50k reads for all methods, but also in this case *mobileRNA* method performed remarkably well with a false positive rate below 5% for almost all conditions. The only exception occurs in the Col-0/C24 sample spiked with 1K reads were about 70% of false positives are retrieved, indicating the if the genomes are similar, lowly expressed mobile transcripts can be more easily confused with noise. All other implemented methods generally performed worst, with a larger number of false positive identified. However, compared to *mobileRNA*, the Wang method retrieved less false positives in the Col-0/C24 sample spiked with 1K of reads, likely due to the higher efficiency of BLAST in identifying the correct location of fewer reads in similar genomes (Figure 3B). Nonetheless, then the false negative rate was calculated, we found that *mobileRNA* outperform all other methods applied to Eggplant/Tomato combination, and all but the Li method for the Col-0/C24 combination. However, the Li method has a false negative rate several order of magnitude higher than *mobileRNA*, generating higher level of noise.

In genomics studies it is not only important to determine if a transcript is mobile, but also the estimation of its accumulation for quantitative comparison studies. However, the noise generated in heterografted analysis could lead to incorrect estimations of the accumulation of mobile transcripts. Therefore, for each method we estimated the accumulation of the simulated mobile transcripts, and we compared it with the expression level (FPKM) calculated for the same genes analysed in their original RNAseq libraries (assumed as expected values of expression). Remarkably, *mobileRNA* displayed a high correlation (Eggplant/Tomato R^2^ = 0.59; Col-0/Ler-o R^2^ = 0.012) in all analysed conditions, in comparison to all other methods (Eggplant/Tomato R^2^ < 0.35; Col-0/Ler-o R^2^ < 0.0.00043). Taken together, these results indicate that mapping RNA samples from heterografted samples on a merged reference strongly reduce the noise generated by false positives and lead to a better estimation of the real accumulation in the plant tissues.

## 4. Discussion

We assembled *mobileRNA,* a user-friendly Bioconductor/R package offering a pre-processing and analysis pipeline to identify the expression of sRNA or mRNA from multiple genomes within a condition. While *mobileRNA* can be used as R interface for standard RNAseq and sRNAseq analysis, it has been specifically develop to detect mobile RNA molecules in grafting experiments. Our approach is based on the use of a merged reference in the alignment step, to find the most probable location of each read during the mapping step, rather than determine them with post-processing analyses. In recent years, similar approaches have been proved valid in population genetic analyses; for example in pan-genome alignment, when references from multiple genomes or transcriptomes are simultaneously used for mapping transcripts within a species, accounting for variations originated from the specific strains/accessions/ecotypes sampled (Bayer, Golicz et al. 2020). Likewise, metagenomic studies aiming to use expression responses sample to explore the diversity of a community in their environment, also benefit from the simultaneous alignment of RNA reads to a database of references (Blanco-Míguez, Beghini et al. 2023, Nam, Do et al. 2023). The use of a merged genome has been also used for the alignment of RNAseq data obtained from tomato hybrids, allowing to discriminate expression originated from different genetic alleles (Lopez-Gomollon, Müller et al. 2022). In our work however, we show that this approach works also when the contribution of the two genomes is not equal, with mobile molecules coming from the distant genome being only a small portion of the total sequenced molecules.

The identification of endogenous mobile RNA molecules in grafting experiments has been associated with a high level of noise, and when potential mobile RNA transcripts identified with genomic approach have been tested with target molecular biology methods, only a small fraction have been validated (Notaguchi, Higashiyama et al. 2015, Kehr, Morris et al. 2022). Recently, a constitutive re-analysis of several dataset sampled from grafted plants has highlighted how the technological and biological noise are responsible for the majority of previously identified mobile transcripts (Paajanen, Tomkins et al. 2024), indicating that more rigorous approaches are necessary for these analyses. Here, we designed a formal approach to evaluate the performances of different mobile RNA detection systems, based on the use of blended datasets and the recovery of false positives. We used this approach to demonstrate that the *mobileRNA* approach outperforms other methods in detecting mobile RNA molecules, substantially reducing the background data noise and increasing the confidence of both mobile sRNA and mobile transcripts. This will allow users to draw more conclusive biological implications from the data and offers a more reliable position to base subsequent analyses and hypotheses. The efficiency in the detection decreases if the two genomes are too similar or the coverage too low (as seen in Arabidopsis accessions, Figure 3). This suggests that the standard coverage of a RNAseq experiments might not be optimal for the detection of lowly accumulated mobile transcripts, and that the use of intra-species grafts could underestimate the number of mobile molecules, compared to the use of inter-species combinations. Nonetheless, the similarity of two genomes is variable across the genome also in distantly related species, and we propose the use of the mappability score (Pockrandt, Alzamel et al. 2020) to estimate the proportion of molecules that could be identified in the merged reference. Our results indicate that, in processing the same grafting combinations of Arabidopsis accessions, the merging of long read-assisted *de novo* assemblies produced significant better mappability scores compared to SNPs-only based assemblies (Figure 2), suggesting that the quality of the reference genomes is also of primary importance for the detection of mobile RNA molecules.

In conclusion, we proposed a new approach for the identification of mobile RNA molecules based on multi-references alignment, characterised by a simple implementation, low data noise and high detection efficiency.

