## Supplementary Figures and Tables for "*mobileRNA*: a tool for efficient analysis of mobile RNA molecules in multiple genomes"



| Supplementary Tables/Figures | Description | Page(s) |
|---|---|---|
| Supplementary Figure 1 | Advanced mobileRNA workflow diagram | 2 |
| Supplementary Figure 2 | Distribution of sequencing reads associated to simulated graft-mobile mRNA transcripts (A) and their associated changes in genome mappability when utilising a merged genome (B). | 3 |
| Supplementary Figure 3 | The distribution of simulated graft-mobile sRNA clusters in abundance and size/length in nucleotides for each analysis produced either by the Random or Tier format. | 4 |
| Supplementary Figure 4 | The distribution of read counts associated with false negatives simulated graft-mobile sRNA clusters. | 5 |
| Supplementary Figure 5 | Comparative analysis of mobile mRNAs identified by the analytical methods in each simulated dataset from 1k to 50k library sizes. | 6 |
| Supplementary Table 1 | The genomes utilised across the analysis | 7 |
| Supplementary Table 2 | Sequencing datasets used in this study for the generation of simulated data. | 8 |
| Supplementary Table 3 | Selected parameters to isolate small RNA clusters from biological samples to be spiked and act as simulated graft-mobile sRNAs for the Random and Tier formats for each analysis | 9 |
| Supplementary Table 4 | Summary statistics of simulated mRNAseq samples | 9 |
| Supplementary Table 5 | Summary statistics of simulated sRNAseq samples | 10 |
| Supplementary Table 6 | Results representing the false positives and false negatives values in the analysis of mRNA simulated datasets. | 11 |
| Supplementary Table 7 | Results representing the false positives and false negatives values in the analysis of sRNA simulated datasets | 12 |

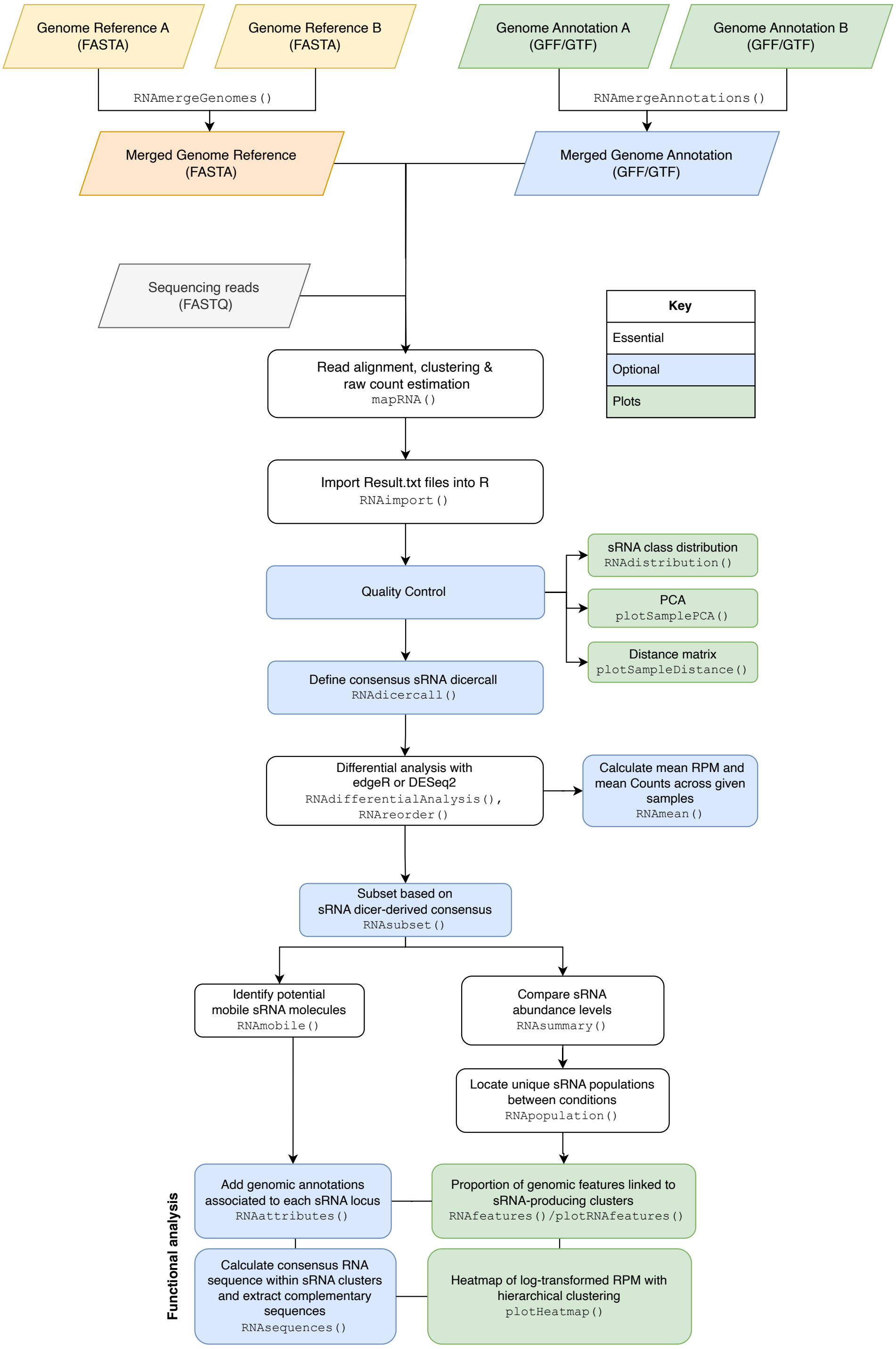


**Supplementary Figure 1. Advanced mobileRNA workflow diagram.** The genome references for the scion and rootstock (represented as Genome Reference A and B) are merged to form a merged genome reference where sequences from each genome remain distinguishable. This is similarly repeated for the genome annotation files to generate a merged genome annotation. The sequencing reads, either from sRNAseq or mRNAseq, are aligned to the merged reference genome, and is followed by clustering or raw count estimation is undertaken. The results are saved to the users desired directory, and then imported into R for downstream analysis, including the selection of putative graft-mobile RNA candidates. Note that, if necessary, more than one distal genome (>2) can be used for complex multi-genome analysis.

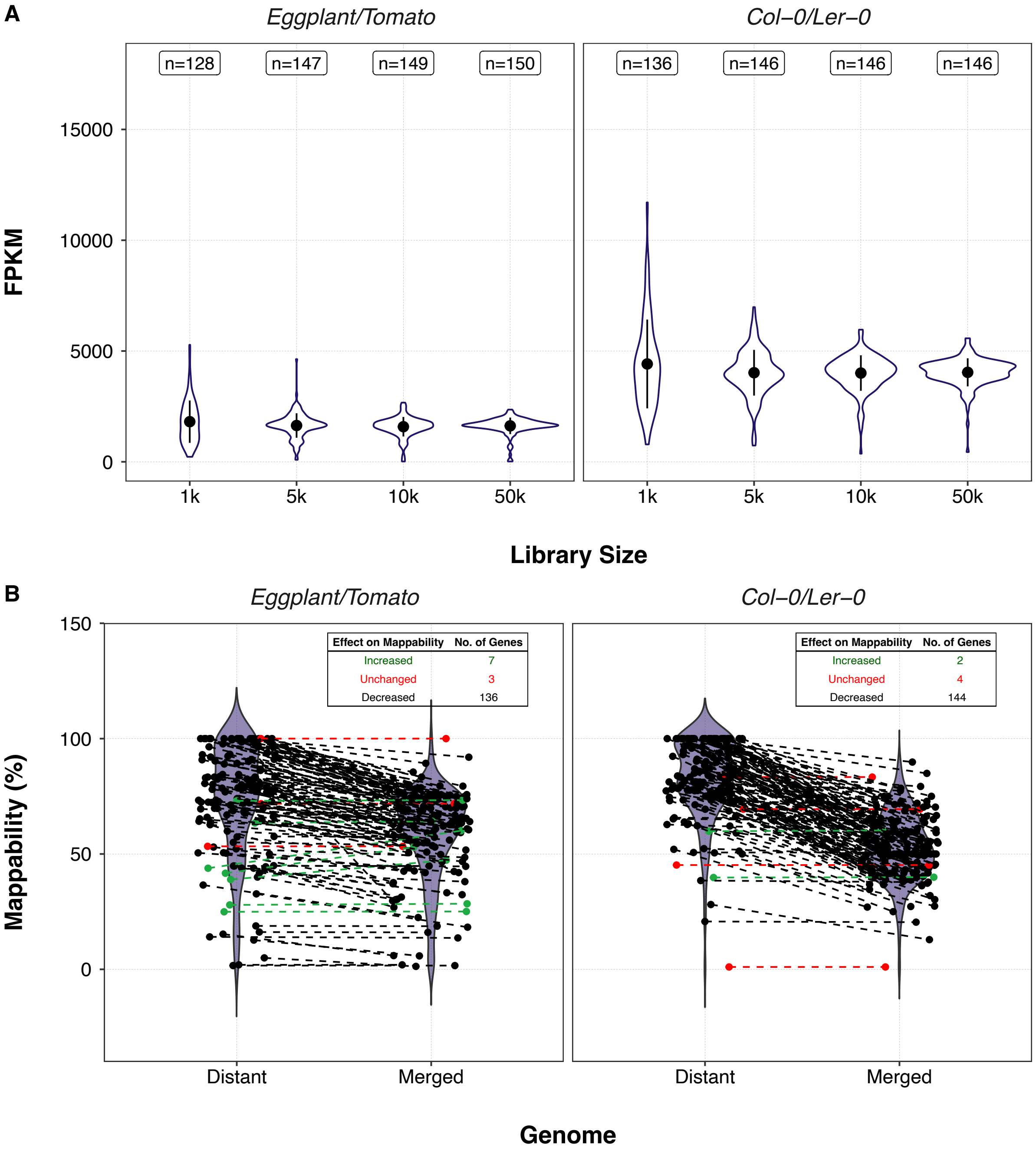


**Supplementary Figure 2. Distribution of sequencing reads associated to simulated graft-mobile mRNA transcripts (A) and their associated changes in genome mappability when utilising a merged genome (B).** A). The distribution of Fragments Per Kilobase of transcript per Million (FPKM) mapped reads of simulated graft-mobile mRNA transcripts of each library size in the Eggplant/Tomato, and Col-0/Ler-0 simulated datasets. B). Genome mappability represents a measure of regions in the genome which are unique or repetitive. Here this is represented as a percentage where 0% refers to regions which lack unique sequences and are highly repetitive while, 100% represents regions which are highly unique. Each point represents a simulated graft-mobile mRNA transcript within their genome of origin (Distant genome) and within the merged genome for the respective analysis. The dash lines connect the genome mappability of the transcript in the distant to the merged genome to illustrate the change in the uniqueness.

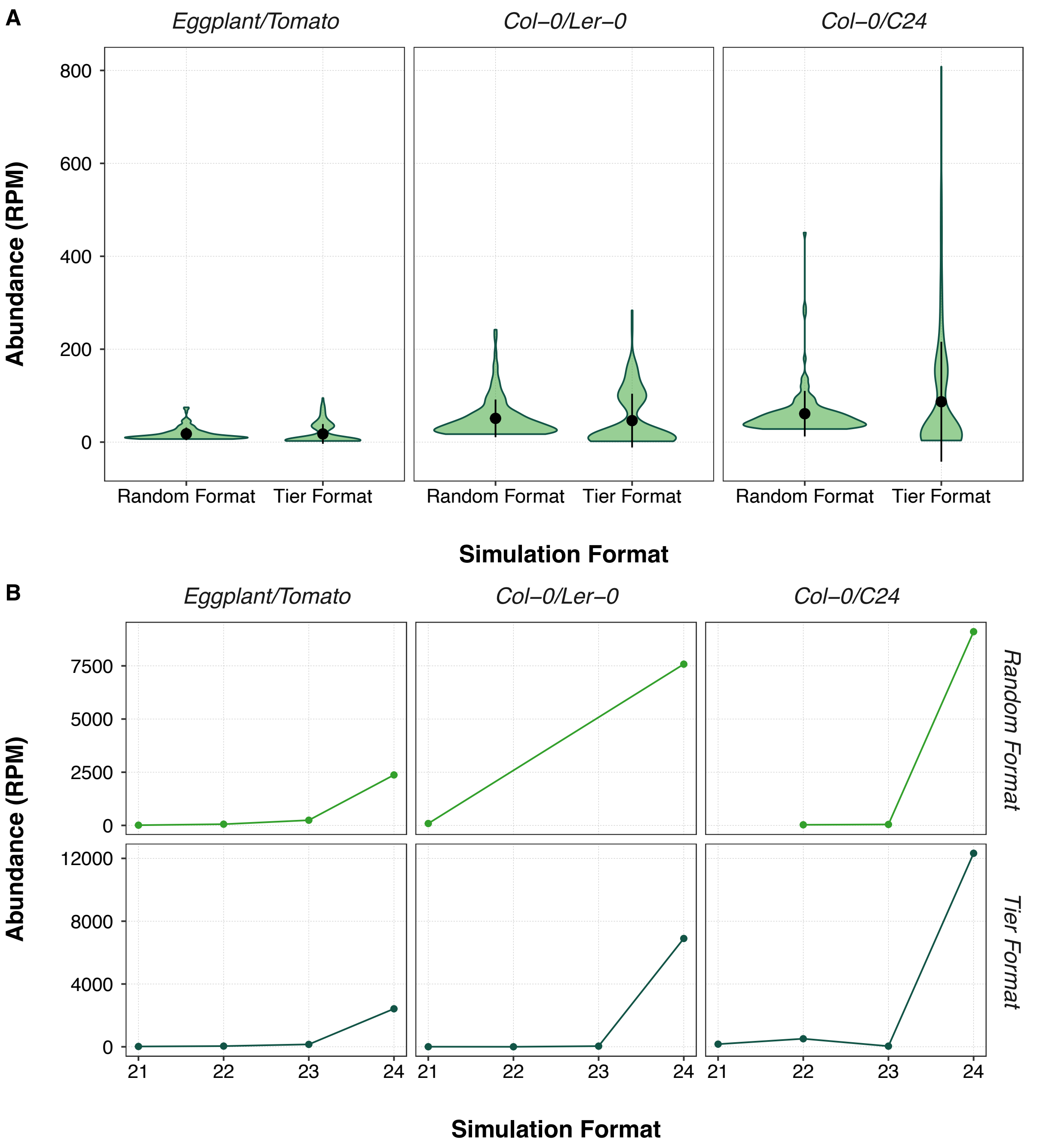


**Supplementary Figure 3. The distribution of simulated graft-mobile sRNA clusters in abundance and size/length in nucleotides for each analysis produced either by the Random or Tier format**. Simulated sRNA clusters were generated by different method including the Random and Tier formats (described in Methods section). A) The distribution in Reads Per Million (RPM) of simulated graft-mobile sRNA clusters B). The distribution in abundance in RPM of sRNA lengths in nucleotides (nt) of simulated graft-mobile sRNA clusters.

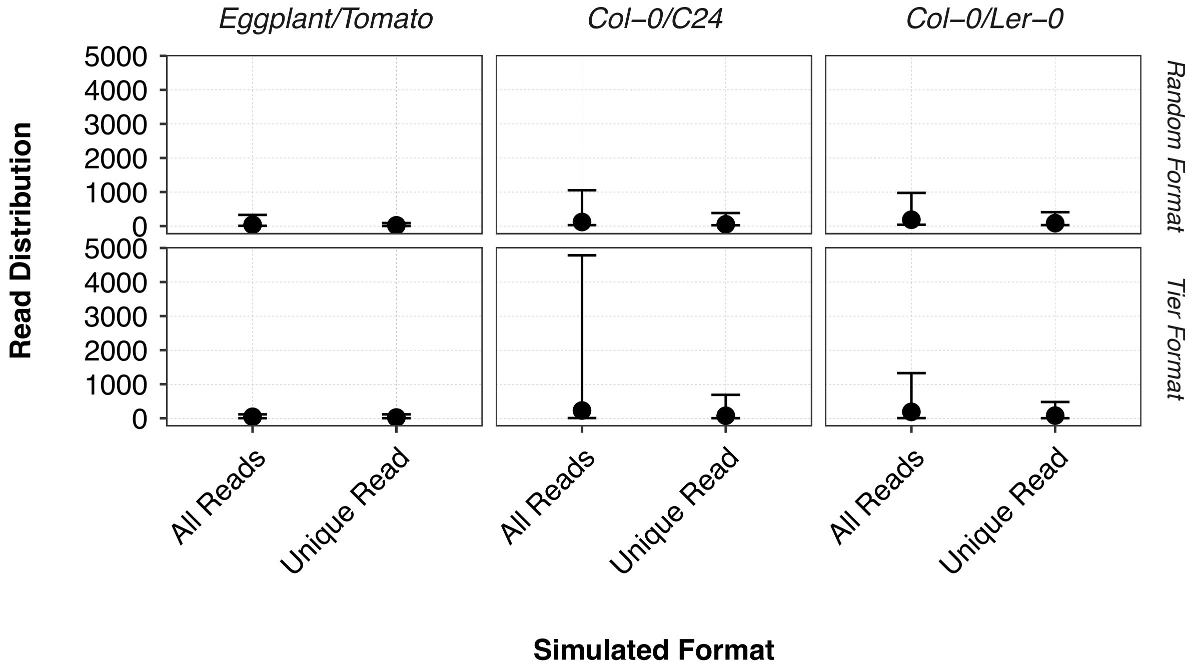


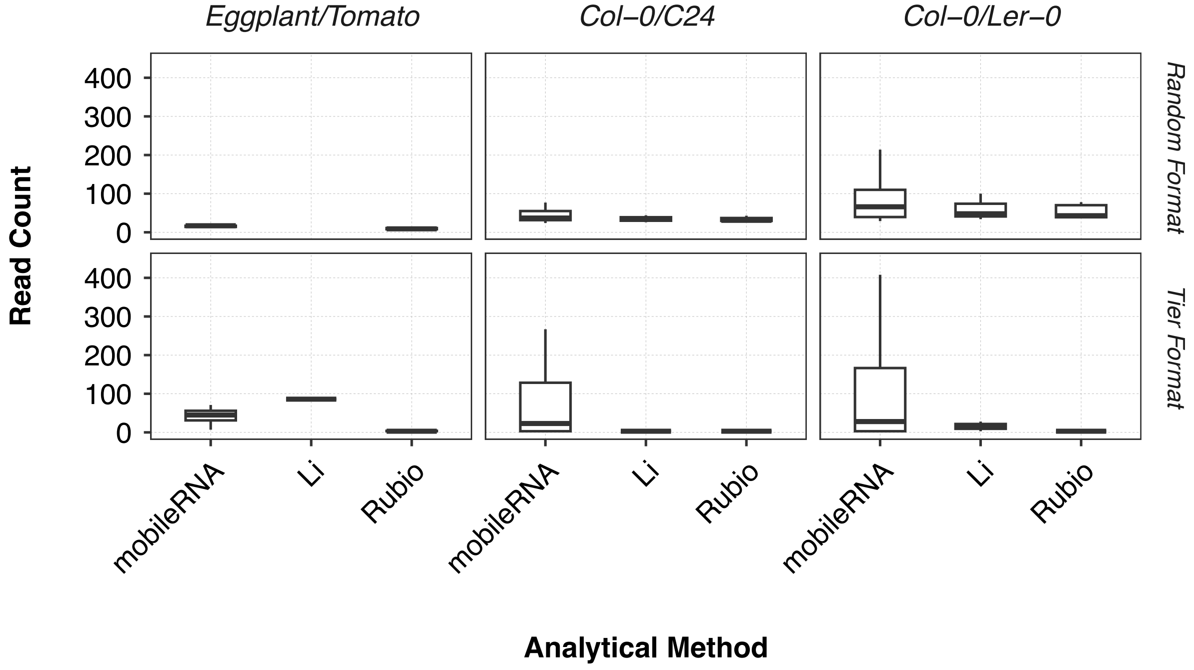


**Supplementary Figure 4. The distribution of read counts associated with false negatives simulated graft-mobile sRNA clusters.** Simulated datasets were produced for three different analyse: Eggplant/Tomato, Col-0/C24, and Col-0/Ler-0 (described in Methods section). A). The distribution in raw read count of simulated sRNA clusters in each analyse for all reads including unique and multimappers in comparison to only the unique reads (reads which aligned to only one location within the Distant genome). B). False negatives refer to the simulated mobile sRNA which were not identified in the analysis. Simulated datasets were analysed by three methods: mobileRNA, Li, and Rubio (described in Methods section). For each method the false negatives were identified (see Supplementary Table 4) and their distribution of reads shown here.

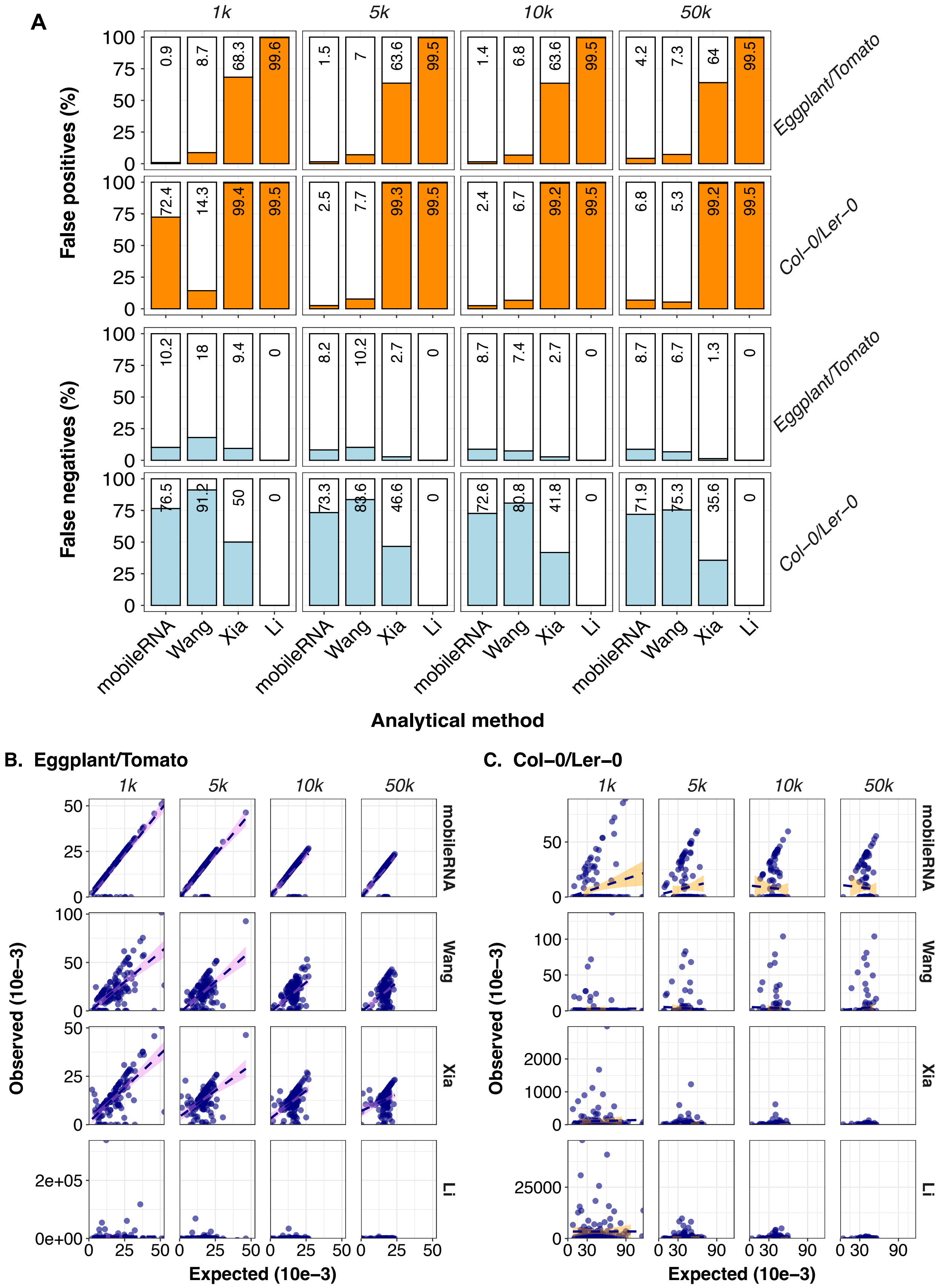


**Supplementary Figure 5. Comparative analysis of mobile mRNAs identified by the analytical methods in each simulated dataset from 1k to 50k library sizes.** The simulated mRNA datasets generate based on different library sizes (1k, 5k, 10k, 50k) were analysed using the four methods: *mobileRNA*, Wang, Xia and Li. True positives (white) refer to the number of mobile RNA correctly identified. False positives (orange) refer to the number of mobile RNA incorrect identified and false negatives (light-blue) refer to the simulated mobile RNA not identified in the analysis. These values are represented as a percentage of the total mobile mRNA located within the dataset. A) Distribution of false positives and false negative to true positives in simulated mRNA datasets. C). Comparison of expected verses observed transcript abundance of simulated mobile mRNAs in the Eggplant/Tomato dataset. The results illustrate a scatter plot for each analytical method (*mobileRNA*, Wang, Xia and Li) displaying the expected verses the observed FPKM values, where each point represents a single simulated transcript. D). Comparison of expected verses observed transcript abundance of simulated mobile mRNAs in the Col-0/Ler-0 dataset. The results illustrate a scatter plot for each analytical method (*mobileRNA*, Wang, Xia and Li) displaying the expected verses the observed FPKM values, where each point represents a single simulated transcript.

**Supplementary Table 1. The genomes utilised across the analysis.** The table displays the source and version of the genome assembly and annotation were extracted from for analysis.

| Organism | Culivar/Accession | Common Name | Genome Assembly & Annotaion | Citation | Size (Mb) |
|---|---|---|---|---|---|
| *Solanum melongena* | '67/3' | Eggplant | Solanum melongena Version 4.1 | Barchi L et al. 2021 | 1,180.0 |
| *Solanumn lycopersicum* | Heinz | Tomato | Solanum lycopersicum Heinz Version 4.0 | Hosmani P et al., 2019 | 795.6 |
| *Arabidopsis thaliana* | Columbia (Col-0) | Thale cress | TAIR10 | Lamesch P et al., 2012 | 121.7 |
| *Arabidopsis thaliana* | Columbia (Col-0) | Thale cress | TAIR10 | Lamesch P et al., 2012 | 121.7 |
| *Arabidopsis thaliana* | Landsberg (Ler-0) | Thale cress | De novo Ler-0 Version 2 | Jiao W et al., 2019 | 122.3 |
| *Arabidopsis thaliana* | C24 | Thale cress | De novo C24 Version 2 | Jiao W et al., 2019 | 121.3 |
| *Vitis vinifera* | Cabernet Sauvignon | Grapevine | Vitis vinifera (PN40024.v4) | French., 2007 | 500 |
| *Vitis riparia* | Gloire de Montpellier | Grapvine | EGFV_Vit.rip_1.0 (GCF_004353265.1) | Girollet et al., 2019 | 500.1 |
| *Phaseolus vulgaris* | G19833 | Common Bean | PhaVulg1_0 (GCF_000499845.1) | Schmutz et al., 2014 | 521.1 |
| *Glycine max* | Williams 82 | Soybean | Glycine_max_v4.0 | Schmutz et al., 2010 | 978.4 |
| *Arabidopsis thaliana* | Columbia (Col) | Thale cress | TAIR10 | Lamesch P et al., 2012 | 121.7 |
| *Nicotiana benthamiana* | - | Tobacco | Nicotiana benthamiana draft genome sequence v2.6.1 | Bombarely et al., 2012 | 378.75 |

**Supplementary Table 2. Sequencing datasets used in this study for the generation of simulated data.** Accessions taken from NCBI Sequence Read Archive (SRA). Information on the appropriate genome assemblies and annotations for each analysis dataset.

| Sequencing type | Dataset | Replicates | Species | Accession/Variety | Simulated genotype | Tissue | Raw reads | Clean reads | Raw data accession | Citation |
|---|---|---|---|---|---|---|---|---|---|---|
| sRNAseq | Tomato/Eggplant | Eggplant_1 | *Solanum melongena* | Line S63 | Tissue (collection) | Anther | 15,431,011 | 10,432,197 | SRR17301403 | Li B et al., 2022 |
| | | Eggplant_2 | *Solanum melongena* | Line S63 | Tissue (collection) | Anther | 11,725,861 | 8,127,789 | SRR17312491 | Li B et al., 2022 |
| | | Eggplant_3 | *Solanum melongena* | Line S63 | Tissue (collection) | Anther | 13,799,582 | 10,451,052 | SRR17312490 | Li B et al., 2022 |
| | | Tomato | *Solanum lycopersium* | BGV007023 | Distant | Leaf | 3,951,725 | 3,828,653 | SRR16077825 | Qing Y et al., 2022 |
| | Col-0/Ler-0 | Col-0_1 | *Arabidopsis thaliana* | Columbia | Tissue (collection) | Root | 15,832,411 | 15,832,411 | SRR8723396 | Blein T et al., 2020 |
| | | Col-0_2 | *Arabidopsis thaliana* | Columbia | Tissue (collection) | Root | 11,546,857 | 11,546,857 | SRR8723397 | Blein T et al., 2020 |
| | | Col-0_3 | *Arabidopsis thaliana* | Columbia | Tissue (collection) | Root | 13,258,374 | 13,258,374 | SRR8723398 | Blein T et al., 2020 |
| | | Ler-0 | *Arabidopsis thaliana* | Landsberg erecta | Distant | Root | 23,009,703 | 23,009,703 | SRR8723405 | Blein T et al., 2020 |
| | Col-0/C24 | Col-0_1 | *Arabidopsis thaliana* | Columbia | Tissue (collection) | Leaf | 15,644,139 | 13,059,905 | SRR21197493 | Yang F et al., 2023 |
| | | Col-0_2 | *Arabidopsis thaliana* | Columbia | Tissue (collection) | Leaf | 16,023,144 | 13,296,185 | SRR21197492 | Yang F et al., 2023 |
| | | Col-0_3 | *Arabidopsis thaliana* | Columbia | Tissue (collection) | Leaf | 15,982,248 | 13,396,813 | SRR21197491 | Yang F et al., 2023 |
| | | C24 | *Arabidopsis thaliana* | C24 | Distant | Leaf | 31,572,522 | 30,057,858 | SRR11521543 | Zhou H et al., 2020 |
| mRNAseq | Tomato/Eggplant | Eggplant 1 | *Solanum melongena* | Ecavi | Tissue (collection) | Leaf | 53,624,938 | 53,624,938 | SRR10060529 | Villanueva G et al., 2023 |
| | | Eggplant 2 | *Solanum melongena* | Ecavi | Tissue (collection) | Leaf | 50,214,317 | 50,214,317 | SRR10060530 | Villanueva G et al., 2023 |
| | Col-0/Ler-0 | Col-0_1 | *Arabidopsis thaliana* | Columbia | Tissue (collection) | Root | 13,982,591 | 13,982,591 | SRR23623443 | PRJNA938935 NCB database (www.ncbi.nlm.nih.g |
| | | Col-0_2 | *Arabidopsis thaliana* | Columbia | Tissue (collection) | Root | 11,038,436 | 11,038,436 | SRR23623442 | PRJNA938935 NCB database (www.ncbi.nlm.nih.g |
| | | Col-0_3 | *Arabidopsis thaliana* | Columbia | Tissue (collection) | Root | 13,485,527 | 13,485,527 | SRR23623441 | PRJNA938935 NCB database (www.ncbi.nlm.nih.g |
| | Col-0/C24 | Col-0_1 | *Arabidopsis thaliana* | Columbia | Tissue (collection) | Root | 13,982,591 | 13,982,591 | SRR23623443 | PRJNA938935 NCB database (www.ncbi.nlm.nih.g |
| | | Col-0_2 | *Arabidopsis thaliana* | Columbia | Tissue (collection) | Root | 11,038,436 | 11,038,436 | SRR23623442 | PRJNA938935 NCB database (www.ncbi.nlm.nih.g |
| | | Col-0_3 | *Arabidopsis thaliana* | Columbia | Tissue (collection) | Root | 13,485,527 | 13,485,527 | SRR23623441 | PRJNA938935 NCB database (www.ncbi.nlm.nih.g |

**Supplementary Table 3. Selected parameters to isolate small RNA clusters from biological samples to be spiked and act as simulated graft-mobile sRNAs for the Random and Tier formats for each analysis.** The coverage selection represents the value or range of selected read coverage to be included in the spike population. The average percentage of spiked reads (Ave. % of spiked reads) within the simulated sRNAseq samples. For each simulated dataset, the average percentage of spiked reads within the total reads was calculated from the simulated replicates within that dataset.

| Simulation Format | Simulated dataset | Tissue genotype | Distant genotype | Ave. % of spiked reads | Coverage selection | Low coverage selection | Mean coverage selection | High coverage selection |
|---|---|---|---|---|---|---|---|---|
| Random Format | Eggplant/Tomato | Eggplant (Solanum melongena) | Tomato (Solanum lycopersicum) | 0.0621 | >7 | - | - | - |
| | Col-0/Ler-0 | Columbia ecotype (Arabidopsis thaliana) | Landsberg ecotype(Arabidopsis thaliana) | 0.2105 | >28 | - | - | - |
| | Col-0/C24 | Columbia ecotype (Arabidopsis thaliana) | C24 ecotype (Arabidopsis thaliana) | 0.0698 | >14 | - | - | - |
| Tier Format | Eggplant/Tomato | Eggplant (Solanum melongena) | Tomato (Solanum lycopersicum) | 0.0678 | - | 3 | 7 | >35 |
| | Col-0/Ler-0 | Columbia ecotype (Arabidopsis thaliana) | Landsberg ecotype(Arabidopsis thaliana) | 0.2116 | - | 3 | 27-28 | >140 |
| | Col-0/C24 | Columbia ecotype (Arabidopsis thaliana) | C24 ecotype (Arabidopsis thaliana) | 0.0470 | - | 3,4,5 | 14-15 | >30 |

**Supplementary Table 4. Summary statistics of simulated mRNAseq samples.** The datasets were formed by using the R package polyester (Frazee, Jaffe et al. 2024) to generate a set of simulated sequencing reads that correspond to a random set of gene transcripts and their associated abundance. These transcripts are generated from the 'Distant' genome and represent the mobile mRNAs in the datasets when spiked/injected into the biological replicates.

| Simulation format | Simulated dataset | Simulated replicates | Tissue genotype | Distant genotype | Cleaned origin reads | Spiked reads | Total reads | % Spiked reads of total |
|---|---|---|---|---|---|---|---|---|
| 1k | Eggplant/Tomato | EggplantTomato_1 | Eggplant (Solanum melongena) | Tomato (Solanum lycopersicum) | 53,624,938 | 1,000 | 53,625,938 | 0.002 |
| | | EggplantTomato_2 | Eggplant (Solanum melongena) | Tomato (Solanum lycopersicum) | 50,214,317 | 1,000 | 50,215,317 | 0.002 |
| | Col-0/Ler-0 | Col0Ler0_1 | Columbia ecotype (Arabidopsis thaliana) | Landsberg ecotype (Arabidopsis thaliana) | 13,982,591 | 1,000 | 13,983,591 | 0.007 |
| | | Col0Ler0_2 | Columbia ecotype (Arabidopsis thaliana) | Landsberg ecotype (Arabidopsis thaliana) | 11,038,436 | 1,000 | 11,039,436 | 0.009 |
| | | Col0Ler0_3 | Columbia ecotype (Arabidopsis thaliana) | Landsberg ecotype (Arabidopsis thaliana) | 13,485,527 | 1,000 | 13,486,527 | 0.007 |
| 5k | Eggplant/Tomato | EggplantTomato_1 | Eggplant (Solanum melongena) | Tomato (Solanum lycopersicum) | 53,624,938 | 5,000 | 53,629,938 | 0.009 |
| | | EggplantTomato_2 | Eggplant (Solanum melongena) | Tomato (Solanum lycopersicum) | 50,214,317 | 5,000 | 50,219,317 | 0.010 |
| | Col-0/Ler-0 | Col0Ler0_1 | Columbia ecotype (Arabidopsis thaliana) | Landsberg ecotype (Arabidopsis thaliana) | 13,982,591 | 5,000 | 13,987,591 | 0.036 |
| | | Col0Ler0_2 | Columbia ecotype (Arabidopsis thaliana) | Landsberg ecotype (Arabidopsis thaliana) | 11,038,436 | 5,000 | 11,043,436 | 0.045 |
| | | Col0Ler0_3 | Columbia ecotype (Arabidopsis thaliana) | Landsberg ecotype (Arabidopsis thaliana) | 13,485,527 | 5,000 | 13,490,527 | 0.037 |
| 10k | Eggplant/Tomato | EggplantTomato_1 | Eggplant (Solanum melongena) | Tomato (Solanum lycopersicum) | 53,624,938 | 10,000 | 53,634,938 | 0.019 |
| | | EggplantTomato_2 | Eggplant (Solanum melongena) | Tomato (Solanum lycopersicum) | 50,214,317 | 10,000 | 50,224,317 | 0.020 |
| | Col-0/Ler-0 | Col0Ler0_1 | Columbia ecotype (Arabidopsis thaliana) | Landsberg ecotype (Arabidopsis thaliana) | 13,982,591 | 10,000 | 13,992,591 | 0.071 |
| | | Col0Ler0_2 | Columbia ecotype (Arabidopsis thaliana) | Landsberg ecotype (Arabidopsis thaliana) | 11,038,436 | 10,000 | 11,048,436 | 0.091 |
| | | Col0Ler0_3 | Columbia ecotype (Arabidopsis thaliana) | Landsberg ecotype (Arabidopsis thaliana) | 13,485,527 | 10,000 | 13,495,527 | 0.074 |
| 50k | Eggplant/Tomato | EggplantTomato_1 | Eggplant (Solanum melongena) | Tomato (Solanum lycopersicum) | 53,624,938 | 50,000 | 53,674,938 | 0.093 |
| | | EggplantTomato_2 | Eggplant (Solanum melongena) | Tomato (Solanum lycopersicum) | 50,214,317 | 50,000 | 50,264,317 | 0.099 |
| | Col-0/Ler-0 | Col0Ler0_1 | Columbia ecotype (Arabidopsis thaliana) | Landsberg ecotype (Arabidopsis thaliana) | 13,982,591 | 50,000 | 14,032,591 | 0.356 |
| | | Col0Ler0_2 | Columbia ecotype (Arabidopsis thaliana) | Landsberg ecotype (Arabidopsis thaliana) | 11,038,436 | 50,000 | 11,088,436 | 0.451 |
| | | Col0Ler0_3 | Columbia ecotype (Arabidopsis thaliana) | Landsberg ecotype (Arabidopsis thaliana) | 13,485,527 | 50,000 | 13,535,527 | 0.369 |

**Supplementary Table 5. Summary statistics of simulated sRNAseq samples.** The datasets were formed by pulling reads from biological samples which were associated to a randomly chosen set of sRNA clusters based on a set of parameters. These sRNA clusters are generated from biological samples associated to the 'Distant' genome and represent the mobile sRNAs in the datasets when spiked/injected into the biological replicates (tissue genotype).

| Simulation Format | Simulated dataset | Simulated replicates | Tissue genotype | Distant genotype | Cleaned origin reads | Spiked reads | Total reads | % Spiked reads of total |
|---|---|---|---|---|---|---|---|---|
| Random Format | Eggplant/Tomato | EggplantTomato_1 | Eggplant (Solanum melongena) | Tomato (Solanum lycopersicum) | 10,432,197 | 5,928 | 10,438,125 | 0.057 |
| | | EggplantTomato_2 | Eggplant (Solanum melongena) | Tomato (Solanum lycopersicum) | 8,127,789 | 5,928 | 8,133,717 | 0.073 |
| | | EggplantTomato_3 | Eggplant (Solanum melongena) | Tomato (Solanum lycopersicum) | 10,451,052 | 5,928 | 10,456,980 | 0.057 |
| | Col-0/Ler-0 | Col0Ler0_1 | Columbia (Arabidopsis thaliana) | Landsberg (Arabidopsis thaliana) | 15,832,411 | 28,106 | 15,860,517 | 0.177 |
| | | Col0Ler0_2 | Columbia (Arabidopsis thaliana) | Landsberg (Arabidopsis thaliana) | 11,546,857 | 28,106 | 11,574,963 | 0.243 |
| | | Col0Ler0_3 | Columbia (Arabidopsis thaliana) | Landsberg (Arabidopsis thaliana) | 13,258,374 | 28,106 | 13,286,480 | 0.212 |
| | Col-0/C24 | Col0C24_1 | Columbia (Arabidopsis thaliana) | C24 (Arabidopsis thaliana) | 13,059,905 | 9,248 | 13,069,153 | 0.071 |
| | | Col0C24_2 | Columbia (Arabidopsis thaliana) | C24 (Arabidopsis thaliana) | 13,296,185 | 9,248 | 13,305,433 | 0.070 |
| | | Col0C24_3 | Columbia (Arabidopsis thaliana) | C24 (Arabidopsis thaliana) | 13,396,813 | 9,248 | 13,406,061 | 0.069 |
| Tier Format | Eggplant/Tomato | EggplantTomato_1 | Eggplant (Solanum melongena) | Tomato (Solanum lycopersicum) | 10,432,197 | 6,474 | 10,438,671 | 0.062 |
| | | EggplantTomato_2 | Eggplant (Solanum melongena) | Tomato (Solanum lycopersicum) | 8,127,789 | 6,474 | 8,134,263 | 0.080 |
| | | EggplantTomato_3 | Eggplant (Solanum melongena) | Tomato (Solanum lycopersicum) | 10,451,052 | 6,474 | 10,457,526 | 0.062 |
| | Col-0/Ler-0 | Col0Ler0_1 | Columbia (Arabidopsis thaliana) | Landsberg (Arabidopsis thaliana) | 15,832,411 | 28,251 | 15,860,662 | 0.178 |
| | | Col0Ler0_2 | Columbia (Arabidopsis thaliana) | Landsberg (Arabidopsis thaliana) | 11,546,857 | 28,251 | 11,575,108 | 0.244 |
| | | Col0Ler0_3 | Columbia (Arabidopsis thaliana) | Landsberg (Arabidopsis thaliana) | 13,258,374 | 28,251 | 13,286,625 | 0.213 |
| | Col-0/C24 | Col0C24_1 | Columbia (Arabidopsis thaliana) | C24 (Arabidopsis thaliana) | 13,059,905 | 6,236 | 13,066,141 | 0.048 |
| | | Col0C24_2 | Columbia (Arabidopsis thaliana) | C24 (Arabidopsis thaliana) | 13,296,185 | 6,236 | 13,302,421 | 0.047 |
| | | Col0C24_3 | Columbia (Arabidopsis thaliana) | C24 (Arabidopsis thaliana) | 13,396,813 | 6,236 | 13,403,049 | 0.047 |

**Supplementary Table 6. Results representing the false positives and false negatives values in the analysis of mRNA simulated datasets.** Simulated datasets were analysed by four methods: mobileRNA, Xia, Li, and Wang (described in Methods section). False positives refer to the number of mobile mRNA incorrect identified while false negatives refer to the simulated mobile mRNA which were not identified in the analysis.

| Simulated Dataset | Analytical Method | Simulation Format | True Positives | False Positives | False Negatives |
|---|---|---|---|---|---|
| Eggplant/Tomato | mobileRNA | 1k | 115 (99.138%) | 1 (0.862%) | 13 (10.156%) |
| | | 5k | 135 (98.54%) | 2 (1.46%) | 12 (8.163%) |
| | | 10k | 136 (98.551%) | 2 (1.449%) | 13 (8.725%) |
| | | 50k | 137 (95.804%) | 6 (4.196%) | 13 (8.667%) |
| | Wang | 1k | 105 (91.304%) | 10 (8.696%) | 23 (17.969%) |
| | | 5k | 132 (92.958%) | 10 (7.042%) | 15 (10.204%) |
| | | 10k | 138 (93.243%) | 10 (6.757%) | 11 (7.383%) |
| | | 50k | 140 (92.715%) | 11 (7.285%) | 10 (6.667%) |
| | Xia | 1k | 116 (31.694%) | 250 (68.306%) | 12 (9.375%) |
| | | 5k | 143 (36.387%) | 250 (63.613%) | 4 (2.721%) |
| | | 10k | 145 (36.432%) | 253 (63.568%) | 4 (2.685%) |
| | | 50k | 148 (36.01%) | 263 (63.99%) | 2 (1.333%) |
| | Li | 1k | 128 (0.437%) | 29134 (99.563%) | 0 (0%) |
| | | 5k | 147 (0.502%) | 29119 (99.498%) | 0 (0%) |
| | | 10k | 149 (0.509%) | 29117 (99.491%) | 0 (0%) |
| | | 50k | 150 (0.513%) | 29116 (99.487%) | 0 (0%) |
| Col-0/Ler-0 | mobileRNA | 1k | 32 (27.586%) | 84 (72.414%) | 104 (76.471%) |
| | | 5k | 39 (97.5%) | 1 (2.5%) | 107 (73.288%) |
| | | 10k | 40 (97.561%) | 1 (2.439%) | 106 (72.603%) |
| | | 50k | 41 (93.182%) | 3 (6.818%) | 105 (71.918%) |
| | Wang | 1k | 12 (85.714%) | 2 (14.286%) | 124 (91.176%) |
| | | 5k | 24 (92.308%) | 2 (7.692%) | 122 (83.562%) |
| | | 10k | 28 (93.333%) | 2 (6.667%) | 118 (80.822%) |
| | | 50k | 36 (94.737%) | 2 (5.263%) | 110 (75.342%) |
| | Xia | 1k | 68 (0.616%) | 10963 (99.384%) | 68 (50%) |
| | | 5k | 78 (0.707%) | 10958 (99.293%) | 68 (46.575%) |
| | | 10k | 85 (0.769%) | 10974 (99.231%) | 61 (41.781%) |
| | | 50k | 94 (0.849%) | 10982 (99.151%) | 52 (35.616%) |
| | Li | 1k | 136 (0.509%) | 26577 (99.491%) | 0 (0%) |
| | | 5k | 146 (0.547%) | 26568 (99.453%) | 0 (0%) |
| | | 10k | 146 (0.546%) | 26570 (99.454%) | 0 (0%) |
| | | 50k | 146 (0.546%) | 26571 (99.454%) | 0 (0%) |

**Supplementary Table 7. Results representing the false positives and false negatives values in the analysis of sRNA simulated datasets.** Simulated datasets were analysed by three methods: mobileRNA, Li, and Rubio (described in Methods section). False positives refer to the number of mobile sRNA incorrect identified while false negatives refer to the simulated mobile sRNA which were not identified in the analysis.

| Simulated Dataset | Analytical Method | Simulation Format | True Positives | False Positives | False Negatives |
|---|---|---|---|---|---|
| Eggplant/Tomato | mobileRNA | Random Format | 145 (96.67%) | 0 (0%) | 5 (3.33%) |
| | | Tier Format | 146 (97.33%) | 1 (0.68%) | 4 (2.67%) |
| | Rubio | Random Format | 145 (96.67%) | 134 (48.03%) | 5 (3.33%) |
| | | Tier Format | 115 (76.67%) | 129 (52.87%) | 35 (23.33%) |
| | Li | Random Format | 150 (100%) | 162314 (99.91%) | 0 (0%) |
| | | Tier Format | 149 (99.33%) | 162317 (99.91%) | 1 (0.67%) |
| Col-0/Ler-0 | mobileRNA | Random Format | 36 (24%) | 0 (0%) | 114 (76%) |
| | | Tier Format | 43 (28.67%) | 0 (0%) | 107 (71.33%) |
| | Rubio | Random Format | 144 (96%) | 12813 (98.89%) | 6 (4%) |
| | | Tier Format | 125 (83.33%) | 12818 (99.03%) | 25 (16.67%) |
| | Li | Random Format | 147 (98%) | 77450 (99.81%) | 3 (2%) |
| | | Tier Format | 148 (98.67%) | 77468 (99.81%) | 2 (1.33%) |
| Col-0/C24 | mobileRNA | Random Format | 90 (60%) | 0 (0%) | 60 (40%) |
| | | Tier Format | 91 (60.67%) | 0 (0%) | 59 (39.33%) |
| | Rubio | Random Format | 147 (98%) | 12062 (98.8%) | 3 (2%) |
| | | Tier Format | 140 (93.33%) | 12072 (98.85%) | 10 (6.67%) |
| | Li | Random Format | 147 (98%) | 19800 (99.26%) | 3 (2%) |
| | | Tier Format | 149 (99.33%) | 44617 (99.67%) | 1 (0.67%) |

### Reference.

Frazee, A., A. Jaffe, R. Kirchner and J. Leek (2024). "polyester: Simulate RNA-seq reads." (R package version 1.39.0.).
